# Functional analysis of gene family evolution demonstrates expansion of digestive, immunity and olfactory functions in the black soldier fly (*Hermetia illucens*) lineage

**DOI:** 10.1101/2025.05.30.657015

**Authors:** Wenjun Zhou, Daniele Kunz, Chris D. Jiggins

## Abstract

Structural variants such as chromosomal rearrangements and gene duplications can play an important role in the adaptation and diversification of organisms. Here, we used comparative genomics to study the functional implications of structural variants across two families of flies. We compared the reference genomes of eight Asilidae species and six Stratiomyidae species, including the black soldier fly (*Hermetia illucens*), a species with an ability to convert organic waste into biomass and a recently expanded global range. The genomes of Stratiomyidae are generally larger than Asilidae and contain a higher proportion of transposable elements, many of which are recently expanded. Gene families showing more gene duplications are enriched for life history related functions such as metabolism in Stratiomyidae and proteolysis and longevity in Asilidae. Gene families showing more gene duplications that are specific to *H. illucens* are mostly related to olfactory and immune responses, while across the Stratiomyidae there is enrichment of digestive and metabolic functions such as proteolysis. Together, our results shed light on gene family expansions that have likely played a role in the ecological success of the black soldier fly.

## Introduction

Structural variations (SVs) are large scale variations in sequence (typically >50bp) caused by insertions, deletions, inversions, duplications and sequence changes due to transposable elements (TEs) (Alkan et al. 2011; Berdan et al. 2024). As a major component of variation in the genome, SVs can play an important role in adaptation and speciation. For example, several inversions between *Drosophila pseudoobscura* and *D. persimilis* are associated with hybrid sterility, maintaining reproductive isolation (Noor et al. 2001; Zhang et al. 2021). In the Colorado potato beetle *Leptinotarsa decemlineata*, SVs are related to insecticide resistance and facilitate the rapid adaptation of this pest species into new agricultural environments (Cohen et al. 2023). In Lepidoptera, SVs play a role in adaptive wing pattern polymorphism (Joron et al. 2011; Brien et al. 2023) and the dynamics of gene family expansion and contraction are associated with diet breadth in butterflies (Dort et al. 2024).

Among many types of SVs, gene copy number variation (CNV) which includes the gain and loss of gene copies, is an important source of genetic variation that can contribute to adaptation. Duplication and deletion of preexisting genes can generate considerable genetic variation and is an important source of variation in genomes (Katju and Bergthorsson 2013). Gene duplications can lead to multiple outcomes. On one hand, duplications of a single gene can create functional redundancy and may affect gene dosage (Hahn 2009; Magadum et al. 2013; Kuzmin et al. 2022). Duplicated genes can also either become a pseudogene, or be subfunctionalized, whereby both genes adopt parts of their original function (Magadum et al. 2013). Alternatively, redundancy can also help the newly duplicated genes “escape” from purifying selection and evolve new functions, known as neofunctionalization (Hahn 2009; Magadum et al. 2013; Birchler and Yang 2022; Kuzmin et al. 2022).

Here we focus on SVs and their functional roles in the black soldier fly (*Hermetia illucens*), a Dipteran species of commercial interest due to its ability to convert organic waste into biomass. *Hermetia illucens* belongs to the Stratiomyidae family (soldier flies), which consists of over 2,700 species found around the world (Woodley and Thompson 2001). Stratiomyidae larvae are often found in water or damp substrates such as decaying organic matter. Despite the similar ecology of many Stratiomyidae species, *Hermetia illucens* stands out as the only species that has spread around the globe as a human commensal and is associated with industrial uses in waste treatment (Nguyen et al. 2015; Tomberlin and van Huis 2020; Siddiqui et al. 2022).

Using a comparative genomics approach, we compared chromosome-level reference genomes of six Stratiomyidae species and eight Asilidae species in (robber flies). During the larval stage, Asilidae are often found in soil or other damp substrates such as rotting organic matter similar to those of Stratiomyidae larvae. However, unlike the Stratiomyidae, Asilidae species are long-lived, with a life span ranging from one to three years, while the Stratiomyidae usually only have short life cycles. Compared to Stratiomyidae adults which usually only feed on plant liquids or do not feed at all, Asilidae adults are predators and feed on other insects. On the phylogeny, Asilidae are one of the families in superfamily Asilidae, which is sister clade to the superfamily Stratiomyomorpha that contains Stratiomyidae (Wiegmann et al. 2011), making this an interesting phylogenetic and phenotypic comparison.

We address the following questions: 1) How much variation in genome size and gene number is there between Stratiomyidae and Asilidae? 2) What is the pattern of types and proportion of repetitive elements among Stratiomyidae and Asilidae species? 3) What are the dynamics of gene birth and death across the phylogeny? 4) How is gene family expansion related to species-specific life history and functional variation?

## Materials and Methods

### Obtaining and pre-processing reference genomes

All genome assemblies were downloaded from the NCBI database with their RefSeq assembly numbers (*Hermetia illucens*: GCF_905115235.1; *Stratiomys singularior*: GCA_954870665.1; *Chorisops tibialis*: GCA_963669355.1; *Beris chalybata*: GCA_949128065.1; *Beris morrisii*: GCA_951812415.1; *Microchrysa polita*: GCA_949715475.1; *Machimus rusticus*: GCA_951509405.1; *Machimus atricapillus*: GCA_933228815.1; *Eutolmus rufibarbis*: GCA_963920795.1; *Tolmerus cingulatus*: GCA_959613345.1; *Neoitamus cyanurus*: GCA_947538895.1; *Leptogaster cylindrica*: GCA_963082835.1; *Dioctria rufipes*: GCA_963924295.1; *Dioctria linearis*: GCA_963930735.1; *Drosophila melanogaster*: GCF_000001215.4) before 1^st^ March 2025. Annotations and peptide sequences of the reference genomes (McCulloch et al. 2023; Thomas et al. 2023; Crowley, Garland, et al. 2024; Crowley, Sivell, et al. 2024; Crowley, Akinmusola, et al. 2024a; Crowley, null, et al. 2024; Crowley, Akinmusola, et al. 2024b; Mitchell et al. 2024a; Mitchell et al. 2024b; Nash et al. 2024; Sivell, Sivell, null, et al. 2024; Sivell, McAlister, et al. 2024; Sivell, Sivell, Mitchell, et al. 2024) were obtained from the Darwin Tree of Life Project (https://wellcomeopenresearch.org/treeoflife) through its online data portal (https://www.darwintreeoflife.org/genomes/genome-notes/) except for *Hermetia illucens* and *Drosophila melanogaster* whose annotations and protein sequences were downloaded from their NCBI RefSeq FTP archive.

### Genome quality assessment

To evaluate the completeness of reference genomes assembled via different pipelines, we used BUSCO 5.8.2 (Simão et al. 2015) to summarize genome completeness and assembly quality before actual analysis. All genomes were compared to the Diptera database diptera_odb10 downloaded from BUSCO website (https://busco-data.ezlab.org/v4/data/lineages/diptera_odb10.2019-11-20.tar.gz). Summary plot was generated using script generate_plot.py implemented in BUSCO based on the output text file of each genome.

### Repetitive elements identification

We used Earl Grey (Baril et al. 2024), an automatic pipeline to identify repetitive elements on each reference genome. For each genome, Earl Grey was run with ten iterations of its “BLAST, Extract, Align, Trim” process. Output annotation and library of each genome were used for Kimura distance calculation using script “divergence_calc.py” implemented in Earl Grey.

### Orthogroups identification and genome-wide synteny analysis

To evaluate orthology relationships among coding genes, we used OrthoFinder 2.5.5 (Emms and Kelly 2019) to assign protein coding genes in all selected 14 species into orthogroups. When running OrthoFinder, maximum likelihood trees were inferred using multiple sequence alignments method (argument “-M msa”). Species tree was visualized using FigTree 1.4.4 (http://tree.bio.ed.ac.uk/software/figtree/).

The results from OrthoFinder were then used as the input for GENESPACE 1.2.3 (Lovell et al. 2022) to construct synteny plots.

### Gene family expansion and contraction analysis

A CAFE5 (Mendes et al. 2021) pipeline was used to calculate the gene birth-death parameter lambda (λ). We used a 3-category Gamma model (-k 3) in which every terminal and non-terminal node has its unique lambda value to better estimate difference in evolutionary rates in multiple species. The output files were summarized and visualized using CafePlotter (https://github.com/moshi4/CafePlotter).

### Functional enrichment analysis

Duplicated genes were extracted from the OrthoFinder output and used for functional enrichment analysis in the Database for Annotation, Visualization, and Integrated Discovery (DAVID) (Sherman et al. 2022). For *Hermetia illucens* genes, gene IDs in its annotation were used directly as the input gene list. For orthogroups with duplications on the most recent common ancestor nodes of Stratiomyidae and Asilidae, enrichment analyses were performed using orthologue gene IDs in the *Drosophila melanogaster* annotation.

Outputs from the enrichment analysis were downloaded as text files and visualized with jvenn (Bardou et al. 2014) and ggplot2 (Wickham 2016).

## Results

### Genome size expansion in Stratiomyidae

We first summarized BUSCO scores of all reference genomes to explore consistency between genomes assembled and annotated by different pipelines. After being compared to a total database of 3,285 Dipteran BUSCO genes, the dataset of 14 reference genomes reached an average of 96.20% completeness (SD = 0.0102), showing consistent level of assembling quality across multiple pipelines (Supplementary Figure 1).

On average, Stratiomyidae (mean genome size = 0.721 gigabytes) have larger genomes compared to Asilidae (mean genome size = 0.559 gigabytes) (Table 1, Supplementary Figure 2). However, *Dioctria linearis* and *D. rufipes* from the Asilidae family have the largest genomes of all species in the study. Average gene number of Stratiomyidae (15,972.83) species also exceeds the one of Asilidae (11,714.25) with the exceptions of *Stratiomys singularior* and *Beris chalybata*. Interestingly, despite having a large genome, the two *Dioctria* species do not possess more genes than other Asilidae species in the dataset (Table 1).

**Table 1.** Size information and BUSCO scores of the reference genomes used in this study.

| Family | Species | Size/Gb | Gene number | Complete BUSCO gene number | Fragmented BUSCO gene number | Missing BUSCO gene number |
| --- | --- | --- | --- | --- | --- | --- |
| Asilidae | *Leptogaster cylindrica* | 0.185 | 10,816 | 3,171 | 10 | 104 |
|  | *Neoitamus cyanurus* | 0.345 | 12,046 | 3,184 | 33 | 68 |
|  | *Eutolmus rufibarbis* | 0.270 | 12,655 | 3,191 | 34 | 60 |
|  | *Machimus rusticus* | 0.264 | 11,510 | 3,195 | 31 | 59 |
|  | *Tolmerus cingulatus* | 0.264 | 12,047 | 3,204 | 27 | 54 |
|  | *Machimus atricapillus* | 0.253 | 10,978 | 3,194 | 30 | 61 |
|  | *Dioctria linearis* | 1.521 | 12,074 | 3,170 | 45 | 70 |
|  | *Dioctria rufipes* | 1.369 | 12,218 | 3,179 | 45 | 61 |
| Stratiomyidae | *Stratiomys singularior* | 0.674 | 11,614 | 3,107 | 49 | 129 |
|  | *Microchrysa polita* | 0.798 | 19,601 | 3,105 | 69 | 111 |
|  | *Hermetia illucens* | 0.948 | 14,004 | 3,116 | 67 | 102 |
|  | *Chorisops tibialis* | 0.816 | 22,148 | 3,147 | 70 | 68 |
|  | *Beris morrisii* | 0.578 | 18,031 | 3,157 | 56 | 72 |
|  | *Beris chalybata* | 0.511 | 10,439 | 3,124 | 62 | 99 |

Genome synteny analysis shows many chromosomal rearrangements across the phylogeny (Figure 1, Supplementary Figure 2). Even between species from the same genus there are a large number of inversion, fission and fusion events (Supplementary Table 1).

**Figure 1.**
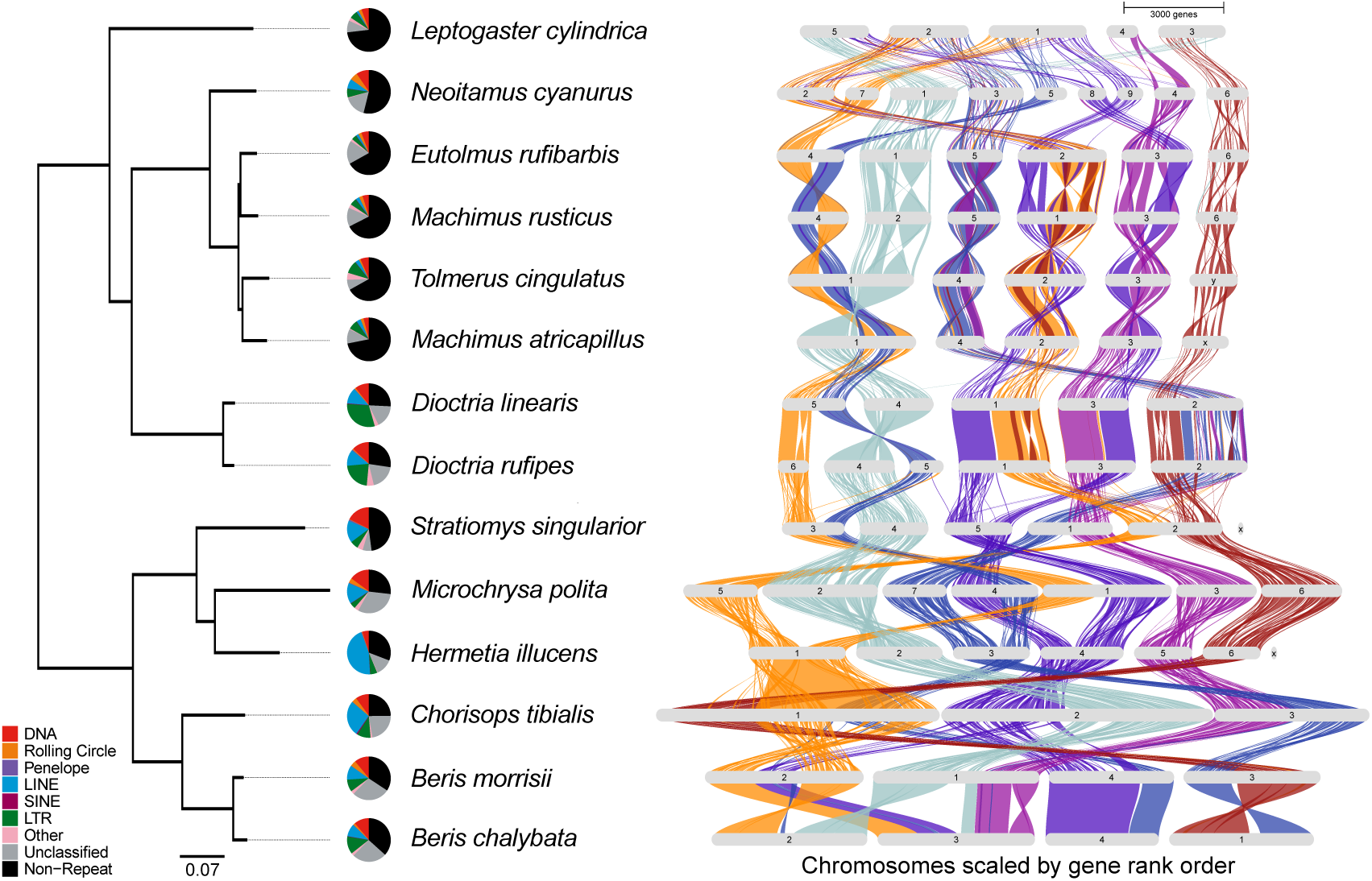
Species tree, proportion of transposable elements (TEs) and genome-wide synteny scaled by physical length of the genomes. Syntenic blocks are divided and aligned based on the order of chromosomes of *Hermetia illucens*. The species tree was rooted using *Drosophila melanogaster*, which is not shown in the tree. Pie charts on the left of species names represent the proportion of different types of transposable elements on the genome. Simple repeats, microsatellites and repeat RNAs were combined as “Other” in the pie charts.

### Variation of repetitive elements across the phylogeny

As expected given their larger genome size, Stratiomyidae have a higher proportion of repetitive elements compared to most Asilidae species except for two *Dioctria* (Figure 1, Supplementary Table 2). There is considerable variation in types of repetitive elements across the phylogeny. For example, long terminal repeats (LTRs) are common in Asilidae genomes, but not in Stratiomyidae. Among Stratiomyidae, DNA repeats dominated in *Stratiomys singularior* and *Microchrysa polita*, but were a relatively low proportion of the repeats in *Hermetia illucens,* where long interspersed nuclear elements (LINEs) were more common compared to other subclasses (Figure 1, Supplementary Table 2).

When divided into subclasses, the majority of repetitive elements show a pattern of recent activity, indicated by the peaks of closely related copies (Supplementary Figure 3). For all species in the dataset, the dominant repetitive elements tend to show lower Kimura 2-Parameter distances (Supplementary Figure 3), showing recent emergence of those repeats. Although the proportion of TEs are generally larger in Stratiomyidae than in Asilidae, the types and divergence time of TEs do not show any strong phylogenetic pattern consistent with very rapid turnover.

### Gene duplication events and gene birth-death rate on terminal and non-terminal nodes

Using *Drosophila melanogaster* as an outgroup, 201,275 out of 211,239 genes (95.3%) were assigned into 15,964 orthogroups, among which 1,707 were species-specific with orthologues found in only one species, and 6,653 had orthologues in all selected species in the dataset (Supplementary Table 3).

Gene duplication events were counted for each of the orthogroups. 32,493 gene duplication events were identified across the phylogenetic tree (Figure 2, Supplementary Table 4). Most of the duplications were on terminal nodes. Compared to Asilidae, Stratiomyidae have more gene duplication events, particularly on terminal nodes. *Stratiomys singularior* has the lowest number of terminal duplication events among the Stratiomyidae species.

**Figure 2.**
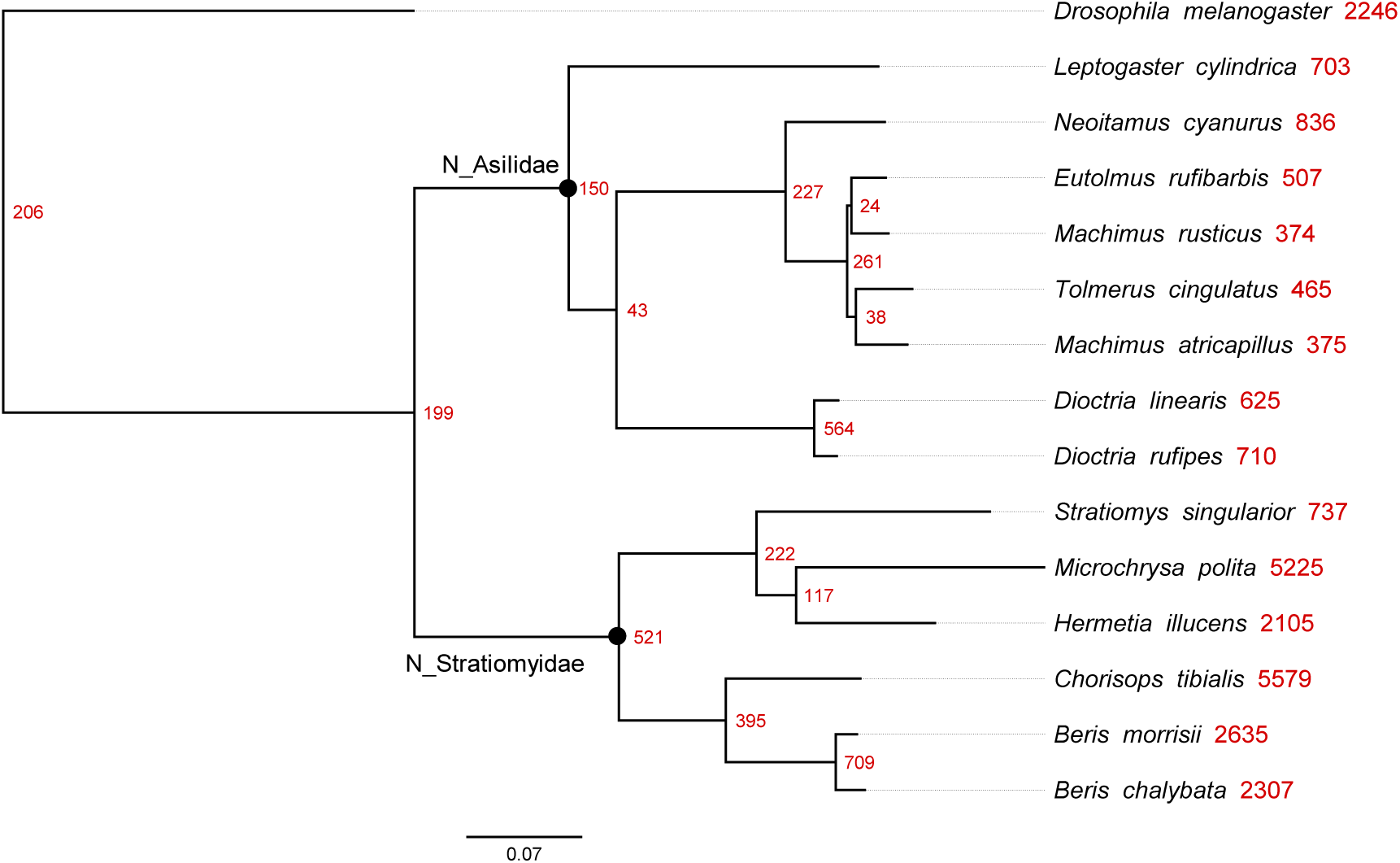
Gene duplication events across the phylogeny. Numbers of duplications on each node are marked on the right side of the node. Only gene duplication events with support higher than 0.5 were considered. The node of the most recent common ancestor of Stratiomyidae and Asilidae were marked as “N_ Stratiomyidae” and “N_Asilidae”.

To study the function of duplicated genes, genes within orthogroups with duplication events showing support values > 0.5 on the most recent common ancestor node of Stratiomyidae (“N_ Stratiomyidae” in Figure 2) and Asilidae (“N_ Asilidae” in Figure 2) were extracted and GO and KEGG enrichment analyses were performed on those genes. In both Stratiomyidae and Asilidae, the enriched biological process term with the highest gene count is “proteolysis” (GO:0006508, *p* = 3.43E-19 for Stratiomyidae and *p* = 1.66E-29 for Asilidae). “Lipid metabolic process” (GO:0006629, *p* = 1.01E-06 for Stratiomyidae and *p* = 1.55E-10 for Asilidae) and “lipid catabolic process” (GO:0016042, *p* = 3.73E-04 for Stratiomyidae and *p* = 4.18E-07 for Asilidae) are also present in both families. Duplicated genes in the common ancestor of Stratiomyidae are mainly enriched for metabolism related biological processes and pathways (Figure 3C, Supplementary Table 5), while duplicated genes in the common ancestor of Asilidae are more enriched for functions related to responses to environmental stimulations and protein refolding (Figure 3D, Supplementary Table 5).

**Figure 3.**
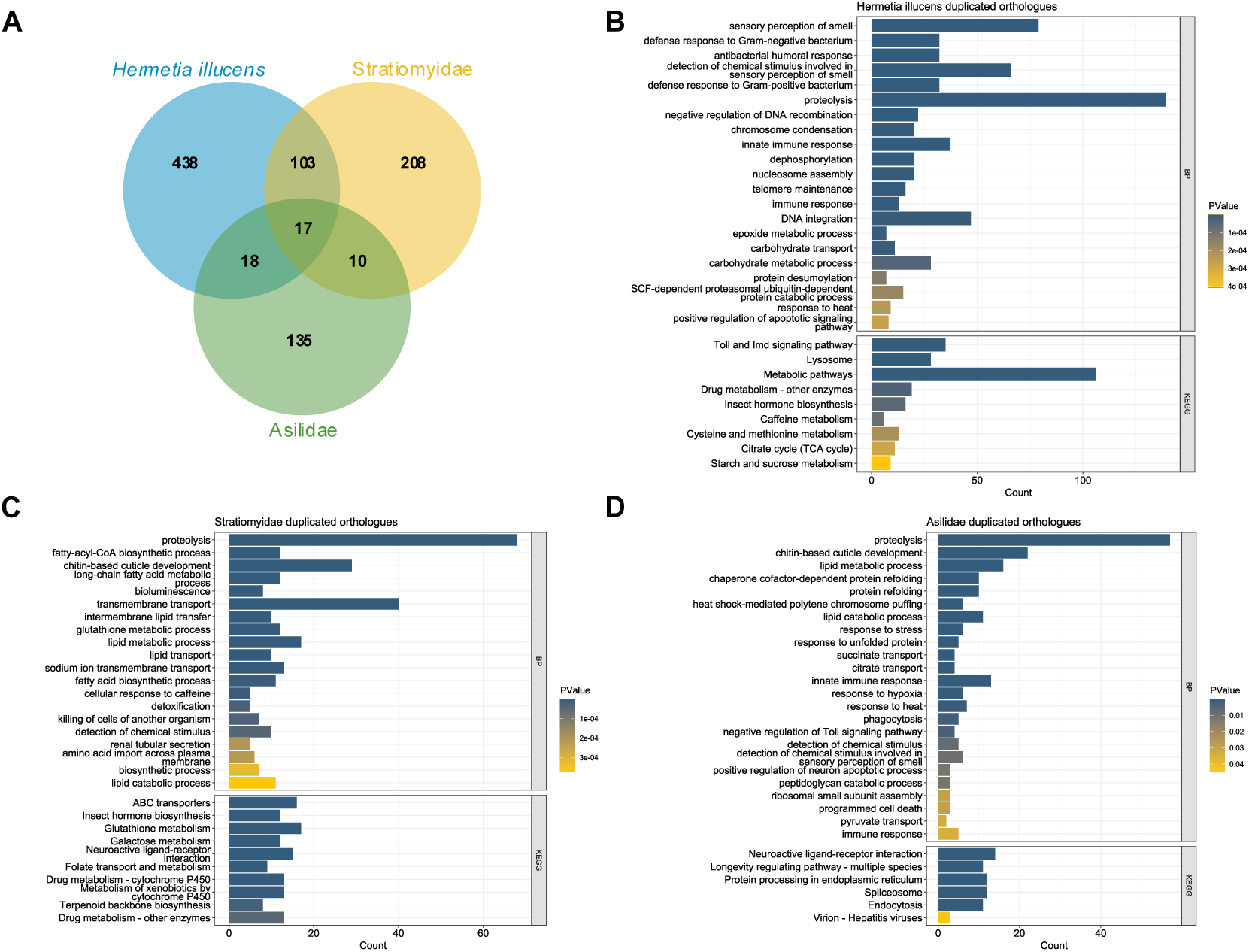
Overlap and functional annotation of duplicated orthogroups among *Hermetia illucens*, Stratiomyidae and Asilidae. **A** Venn diagram of orthogroups with duplication events. **B** GO biological process and KEGG enriched functional terms of *Hermetia illucens* specific duplicated genes. **C** GO biological process and KEGG enriched functional terms of duplicated genes in the most recent common ancestor node of Stratiomyidae. **D** GO biological process and KEGG enriched functional terms of duplicated genes in the most recent common ancestor node of Asilidae.

Although several functional terms showing gene duplications were shared between families, the numbers of overlapping orthogroups was limited. Only 27 orthogroups were shared among the 180 orthogroups that had high support for duplication events in the common ancestor of Asilidae and 338 in Stratiomyidae (Figure 3A). There was greater overlap between orthogroups with duplications in the common ancestor of Stratiomyidae and those in *Hermetia illucens* (Figure 3A). After excluding those orthogroups that are shared, duplicated genes on terminal node of *Hermetia illucens* showed a different pattern compared to those duplicated in common ancestor of Stratiomyidae. Aside from proteolysis which still had the highest number of enriched genes, more genes were enriched in immune and antibacterial responses and olfaction (Figure 3B). In KEGG pathways, the most enriched terms are metabolism-related, while “Toll and Imd signaling pathway”, involved in the immune response, is also enriched (Figure 3B). These terms are consistent with adaptation to the decomposing-related life history of *H. illucens*.

We used the gene birth-death parameter λ, which is the probability of any gene to be gained or lost, as calculated in the CAFE5 pipeline (Mendes et al. 2021) to infer the pattern of expansion and contraction of orthogroups across the phylogeny. Each orthogroup assigned in the previous analysis is considered as a “gene family” and its size was mapped to each node on the species tree. Under a Gamma model with 3 Gamma rate categories, the λ value on each node varies from 0.00002 to 0.00388 (Figure 4). The gene birth-death parameter is higher on terminal nodes compared to non-terminal nodes and reaches the highest on *Neoitamus cyanurus* in Asilidae and *Hermetia illucens* in Stratiomyidae (Figure 4), indicating the highest rate of gene family evolution in these species.

**Figure 4.**
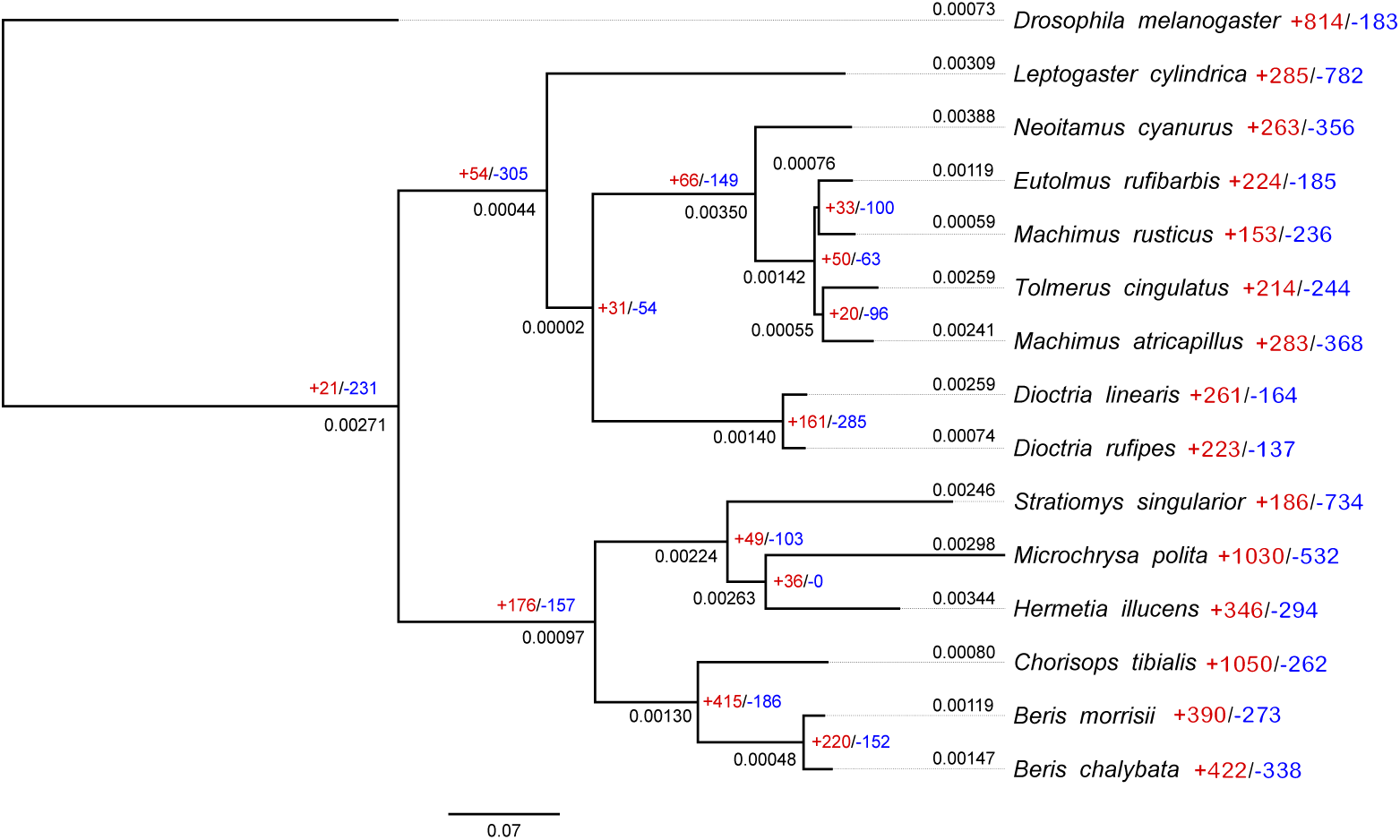
Gene family evolution across the phylogeny. Number on the left side of each node represents the gene birth-death parameter λ of this node. Numbers with “+” and “-” beside each node represent the total number of expanded (“+”) and contracted (“-”) orthogroups on this node.

In *Hermetia illucens*, 109 orthogroups showed significant changes from the last common ancestor node (Supplementary Table 6), most of which are expanded but not contracted. These significantly expanded orthogroups include several gene families involved in immune response (OG0000160 and OG0000220) and olfaction (OG0000020 and OG0000025) (Supplementary Table 6, Supplementary Table 7).

In the top twenty expanded orthogroups in *Hermetia illucens*, only two (OG0000005 and OG0000019) has a lower copy number compared to its closest relative *Microchrysa polita* (Supplementary Table 7). In general, when compared to the most recent common ancestor node of Stratiomyidae, many of these orthogroups expanded in both species, but their expansions in *H. illucens* are larger than in *Microchrysa polita*.

## Discussion

The black soldier fly *Hermetia illucens* has become a major commercial insect species used to consume organic waste. In addition to its value in the industry, it also provides an interesting example of rapid range expansion and human-mediated evolution. By comparing high quality chromosome-level reference genomes from the Stratiomyidae and Asilidae families, we provide evidence for the adaptation of *H. illucens* and its close relatives to their specific life history. We have shown recent activity of transposable elements, as well as large amounts of gene duplication on terminal nodes, high gene birth-death rate and expansions of metabolism and immune related gene families.

Transposable elements have been suggested to be an important factor driving insect evolution (Gilbert et al. 2021). We found varying proportions of transposable elements ranging from 26.85% of the genome in *Leptogaster cylindrica* to 75.18% in *Chorisops tibialis* (Supplementary Table 2), which correspond to previous findings in Diptera (Petersen et al. 2019). The TEs in Stratiomyidae and Asilidae not only vary in abundance but also divergence time. In all species DNA, LINEs and LTRs show recent activity which can be represented by the peaks of base pair numbers with Kimura 2-Parameter Distance near zero (Supplementary Figure 3). This suggests a recent expansion of these TEs in both families. However, in some species, a smaller proportion of other TE superfamilies also show peaks with a much higher Kimura 2-Parameter Distance, such as SINEs in *Leptogaster cylindrica* (Supplementary Figure 11A), *Neoitamus cyanurus* (Supplementary Figure 11B), *Eutolmus rufibarbis* (Supplementary Figure 11C) and *Hermetia illucens* (Supplementary Figure 11K), while the same TE superfamily does not exist in *Dioctria rufipes* (Supplementary Figure 11H), *Stratiomys singularior* (Supplementary Figure 11I), and *Beris morrisii* (Supplementary Figure 11M). In general, the types of TEs in Stratiomyidae and Asilidae correspond to phylogenetic relationships, with similar proportion of TE types are found in more closely related species (Figure 1), which is consistent with previous findings in the majority of insects (Gilbert et al. 2021). However, the distribution of Kimura 2-Parameter Distance of different TEs types does not show a similar phylogenetic pattern, consistent with the patterns found in *Drosophila* (Petersen et al. 2019).

Domestication represents an excellent system in which to study recent adaptation. *Hermetia illucens* is a putative example of recent domestication, with populations brought into captivity repeatedly around the world. Changes of food supply often lead to metabolism-related functional shifts in domesticated animals (Gering et al. 2019). In other species, domestication is associated with changes in diet (Luca et al. 2010), immunity (Chen et al. 2017) and digestive system gene functions (Axelsson et al. 2013; Chen et al. 2017; Glazko et al. 2021). The individual of *Hermetia illucens* that was sequenced for the reference genome was from an industry strain (Generalovic et al. 2021), so we cannot here distinguish between adaptations associated with the species generally and those that have happened since its domestication. It is possible that duplicated genes related to carbohydrate metabolic process that are not shared by other Stratiomyidae species (Figure 3B) are related to the recent domestication of *H. illucens*. In addition to carbohydrate, several other metabolic pathways can also be found in the enriched functional terms, such as drug, cysteine, methionine, starch, sucrose and even caffeine metabolism (Figure 3B). These enriched pathways could be associated with higher efficiency of *H. illucens* on certain diets compared to other Stratiomyidae species, and potential influences of the domestication process. Two major functions, olfactory sensory and immune responses, are specifically enriched in *H. illucens* compared to the most recent common ancestor node of Stratiomyidae (Figure 3A, Table 2). The expansion of olfactory and immune related genes in *H. illucens* can potentially explain the reason that it has become so widely used in bioconversion of organic waste (Tomberlin and van Huis 2020) and is often seen as “dominant” species in compost piles. All these offer clues into how *H. illucens* has adapted to human influenced diets compared to its Stratiomyidae relatives.

Despite the differences in both adult diets, habitats and life span, the top hit in biological process GO terms is proteolysis (Figure 3D) for both Stratiomyidae and Asilidae. This suggests that some specific gene families and their function may have been involved in adaptation to different life history traits. The same pattern can be observed for lipid metabolic process (Figure 3C&D). However, more enriched functional terms in Asilidae are related to responses to external stimuli, including response to stress, heat and hypoxia (Figure 3D). One of the KEGG pathway top hits in Asilidae, longevity regulating pathway, is not enriched for genes that are duplicated in Stratiomyidae, which may be associated with the longer life span of adults in Asilidae compared to Stratiomyidae.

Apart from those gene families that were expanded, several gene families have been completely lost in *H. illucens* (Supplementary Table 6). One example is the gene family which includes *Or83c* and its orthologues (orthogroup OG0000615). In *Drosophila melanogaster*, this odorant receptor is related to sensitivity to farnesol, a fruit rind volatile present in many ripe citrus rinds (Ronderos et al. 2014). The fact that this gene and its orthologues were lost while other groups of olfactory receptor genes such as *Or2a*, *Or59a*, *Or33a*, *Or33b*, *Or33c*, *Obp44a*, *Obp83g*, *Obp99a*, *Obp99b* and *Obp99c* are largely expanded in *H. illucens* suggests potential species-specific diet preference and diversification in olfactory related gene functions.

## Conclusions

Structural variants, gene duplications in particular, are a driving force of functional adaptation. Using a comparative genomics approach, we show evidence of functional adaptation of the soldier flies (Stratiomyidae) and the robber flies (Asilidae) to their habitats and life history. We found a generally larger proportion of transposable elements in Stratiomyidae compared to Asilidae, and the majority of these are recently expanded. More gene duplication events were found on terminal nodes in Stratiomyidae than in Asilidae, and only 27 duplicated orthogroups are shared between the most recent common ancestors of these two families. We found 120 duplicated orthogroups that are shared between the most recent common ancestor of Stratiomyidae and *H. illucens*, indicating functional similarity of gene duplications. Certain genes involved in metabolism-related functions in *Hermetia illucens* are also duplicated in the common ancestor of Stratiomyidae, but more duplicated genes are related to olfactory sensory and immune response in *H. illucens*, which also showed the highest gene birth-death parameter in Stratiomyidae. This suggests a higher rate of gene family evolution and functional specialization in gene families that are beneficial to the decomposing life history in *H. illucens*. Together, our results provide insights into the relationship between structural variation, especially gene duplications, and functional adaptation in two diverse families. We also provide directions for future experimental validation on these gene families and their specific role in functional pathways such as immune response, proteolysis and other metabolism.

## Acknowledgements

We thank Darwin Tree of Life project (https://www.darwintreeoflife.org/) and Wellcome Sanger Institute for the public reference genome resources and the guidance on data portal usage. WZ acknowledges support of the Cambridge Trust and China Scholarship Council (202206380035).

---

**Supplementary Figure 1.**
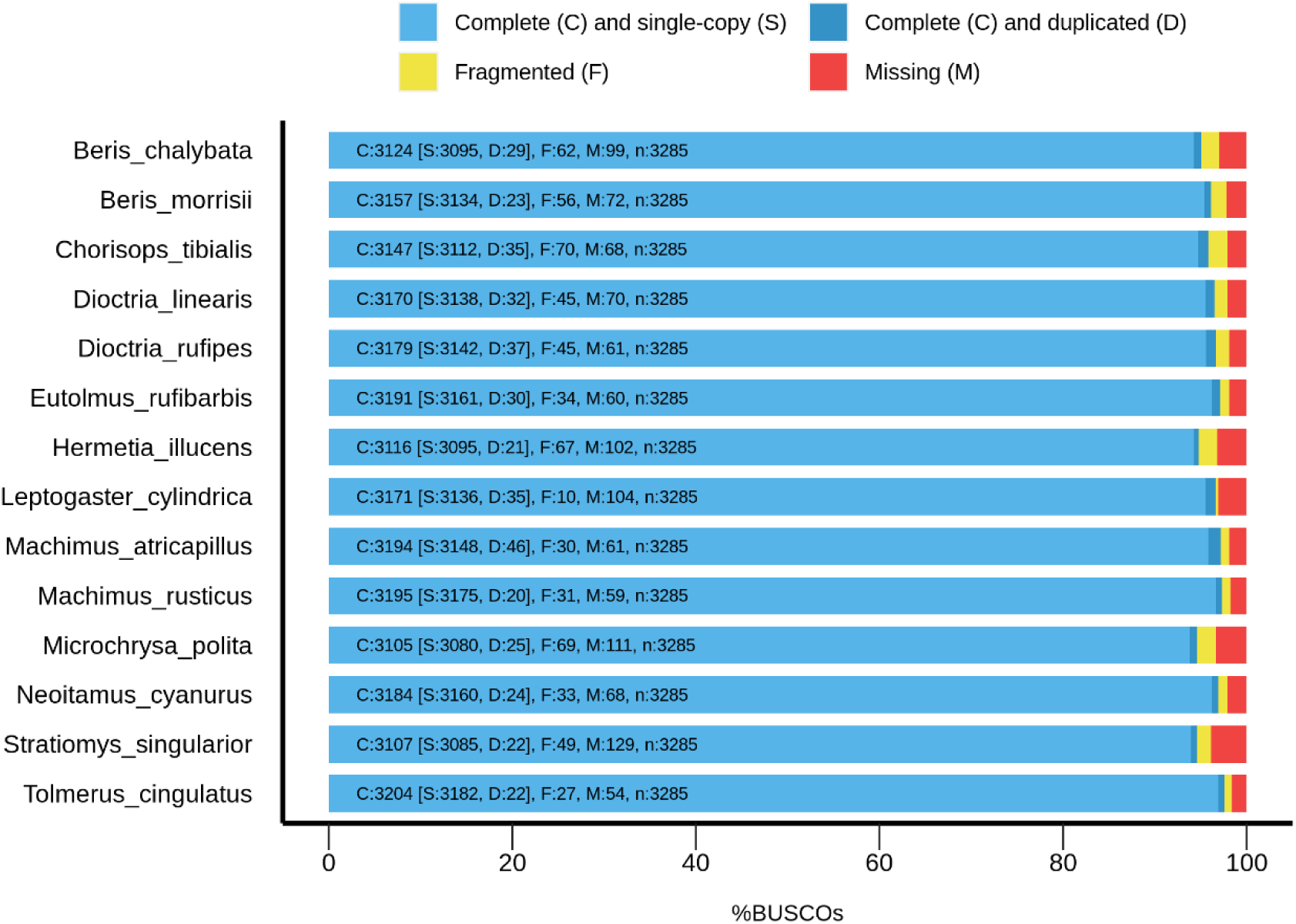
BUSCO assessment summary of all reference genomes used in this study. X axis represents the proportion of BUSCO genes compared to all BUSCO genes in the Dipteran database.

**Supplementary Figure 2.**
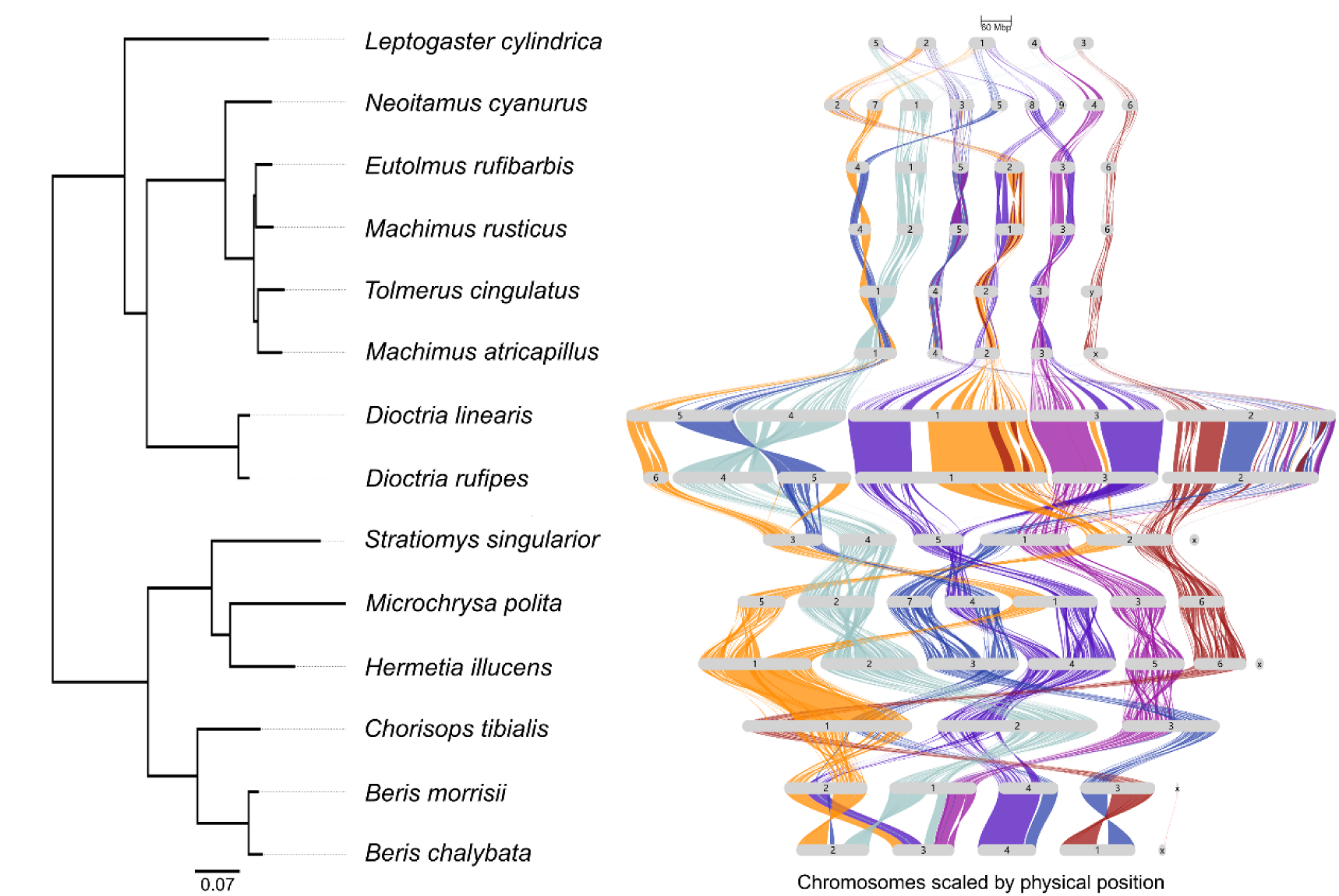
Genome-wide synteny scaled by physical length of the chromosomes. Syntenic blocks are aligned based on the order of chromosomes of *Hermetia illucens*. Species tree was rooted using *Drosophila melanogaster* as outgroup which is not shown in the figure.

**Supplementary Figure 3.**
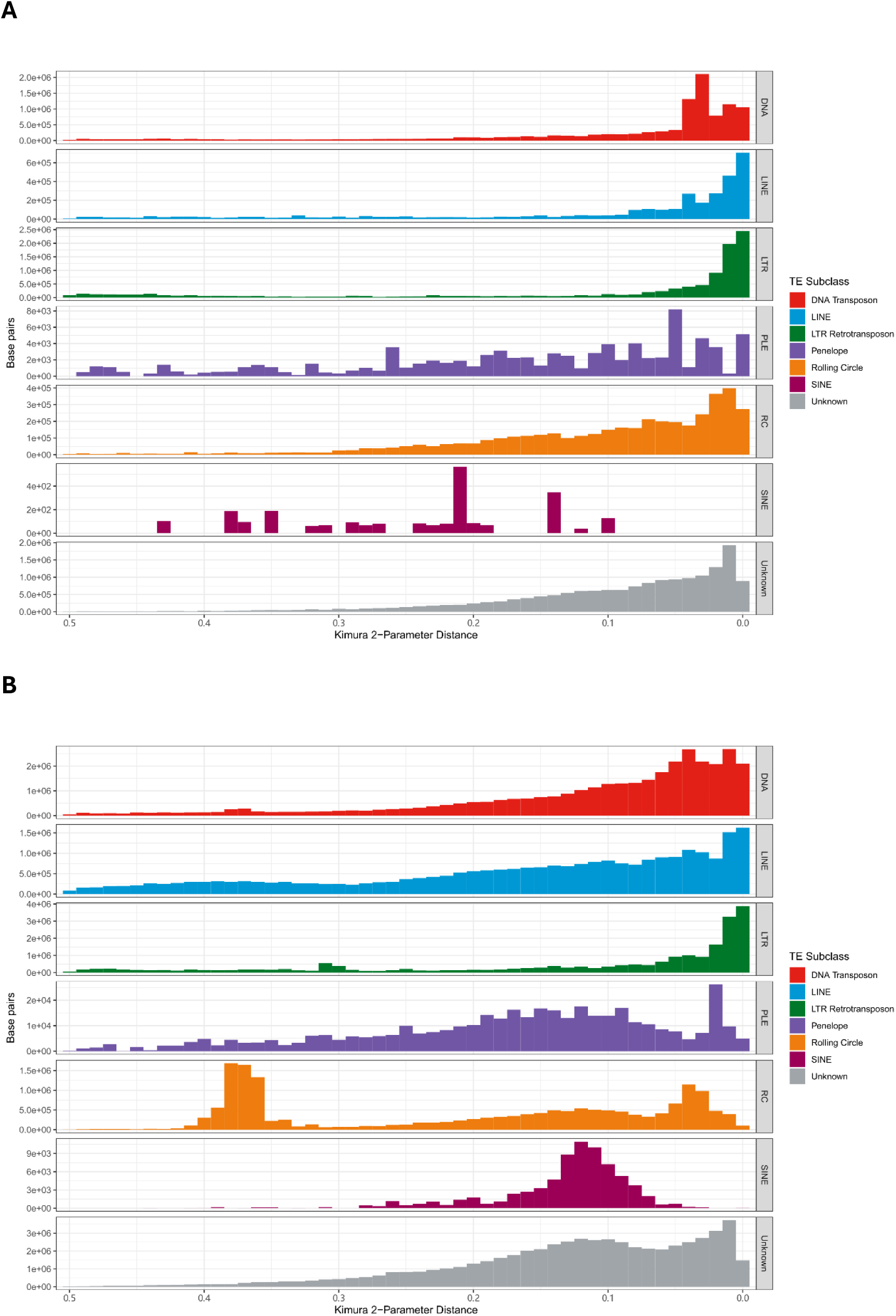
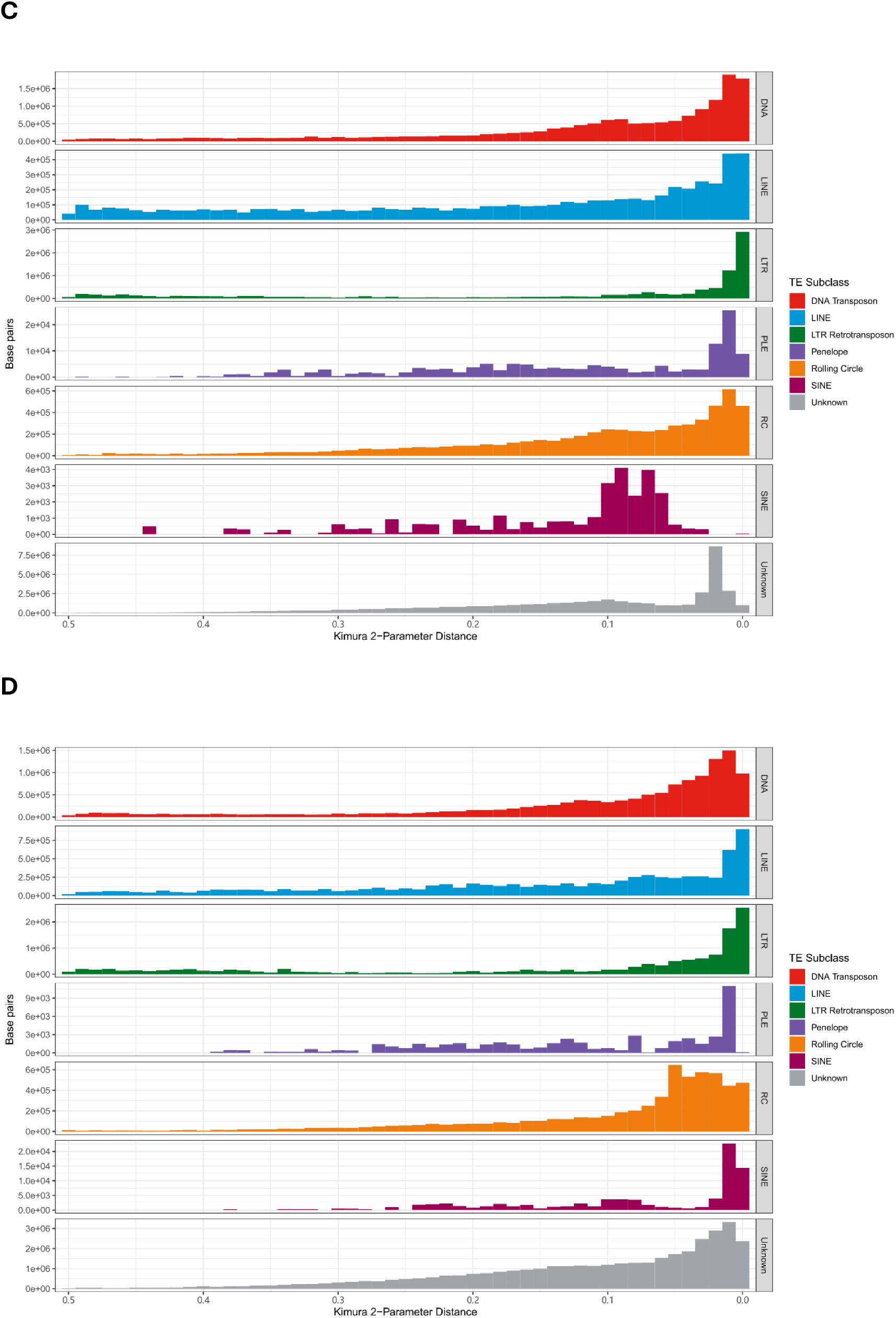
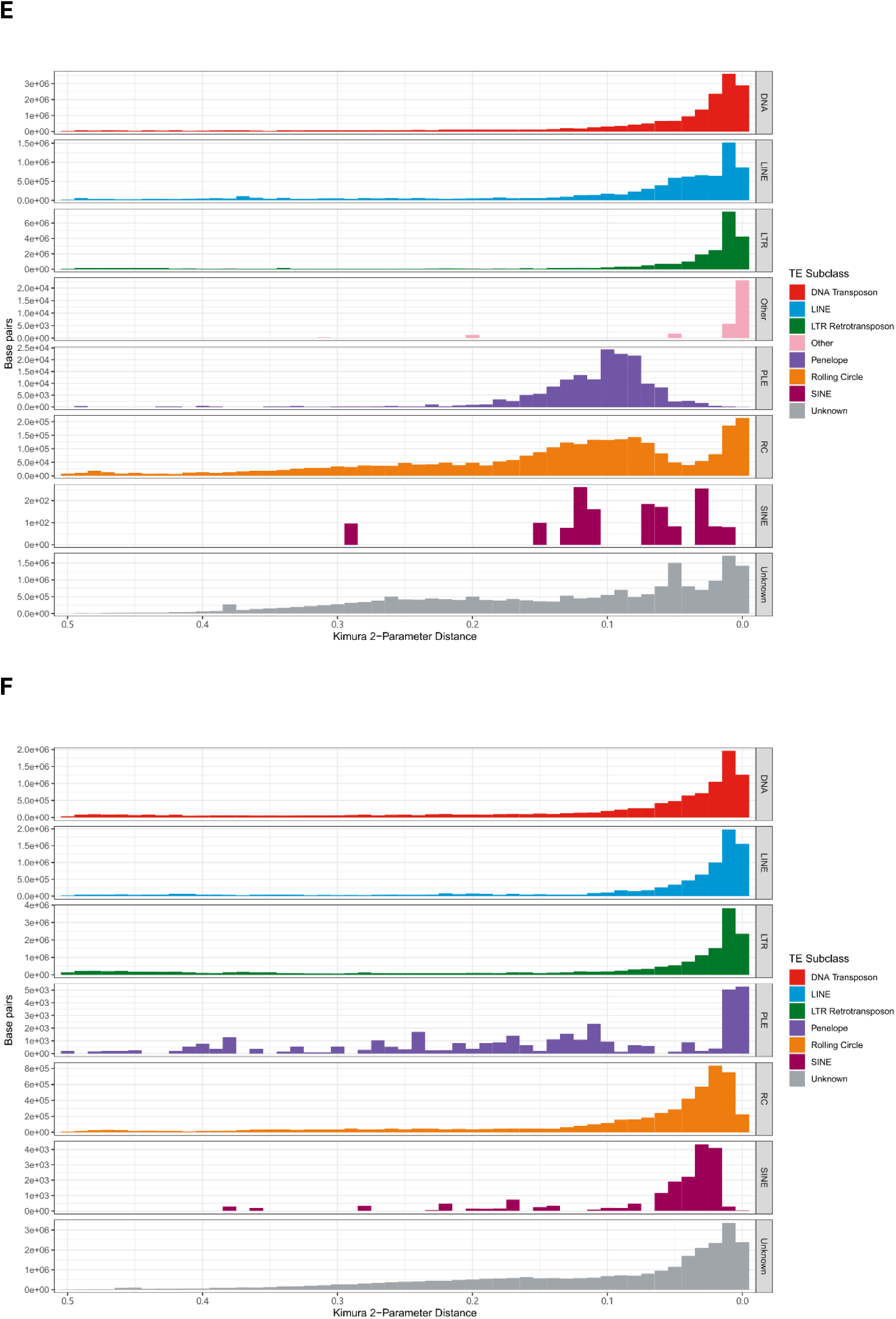
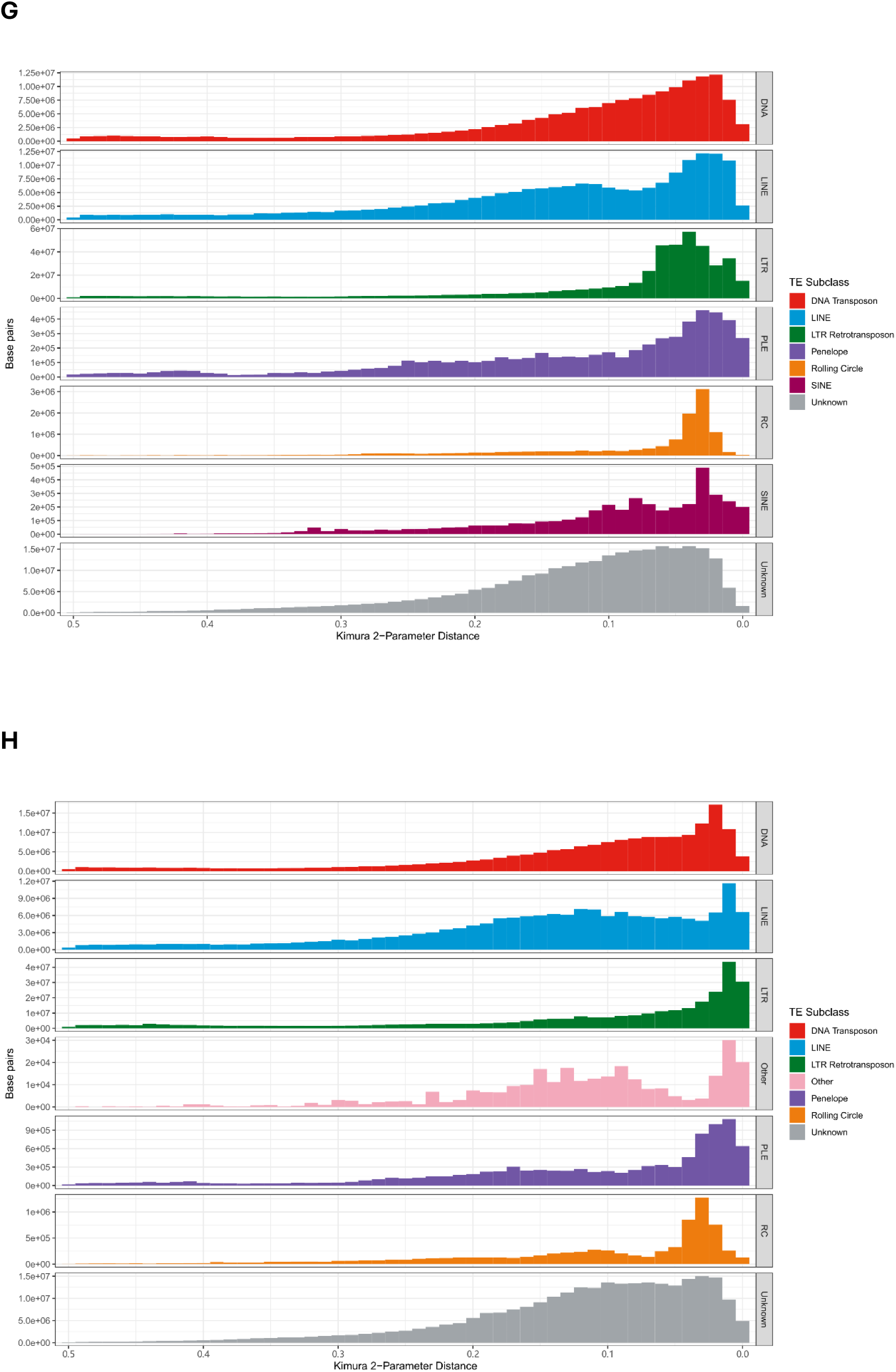
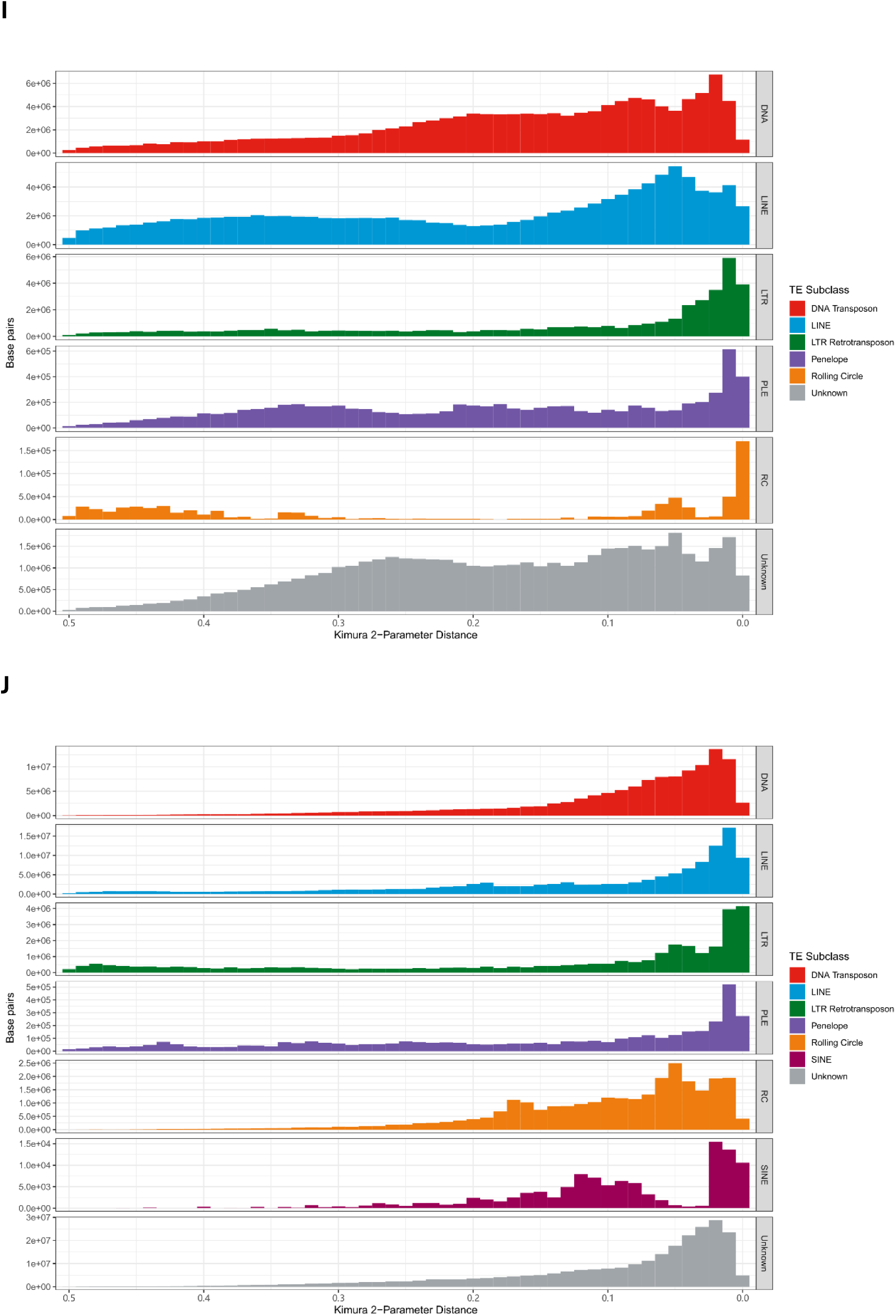
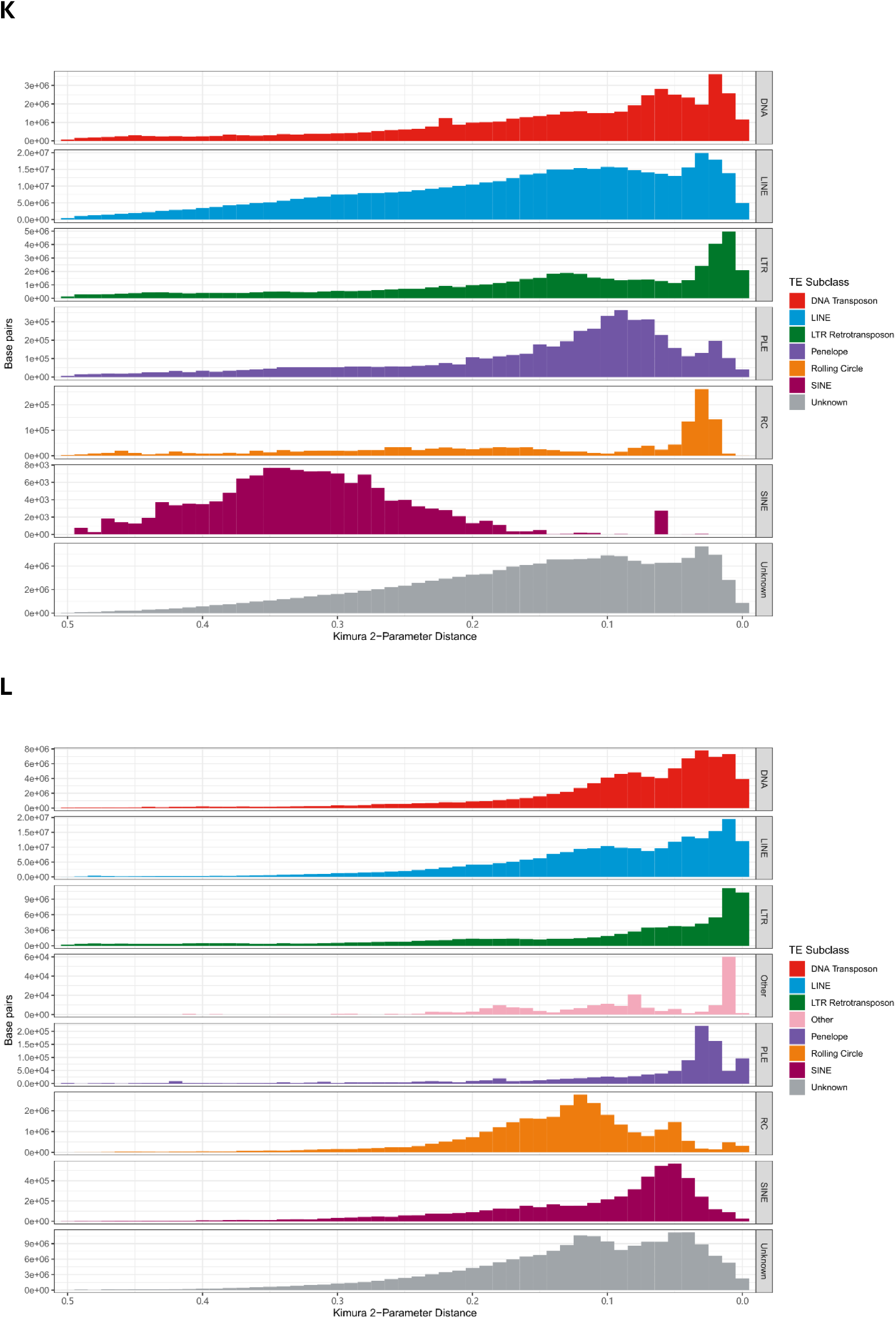
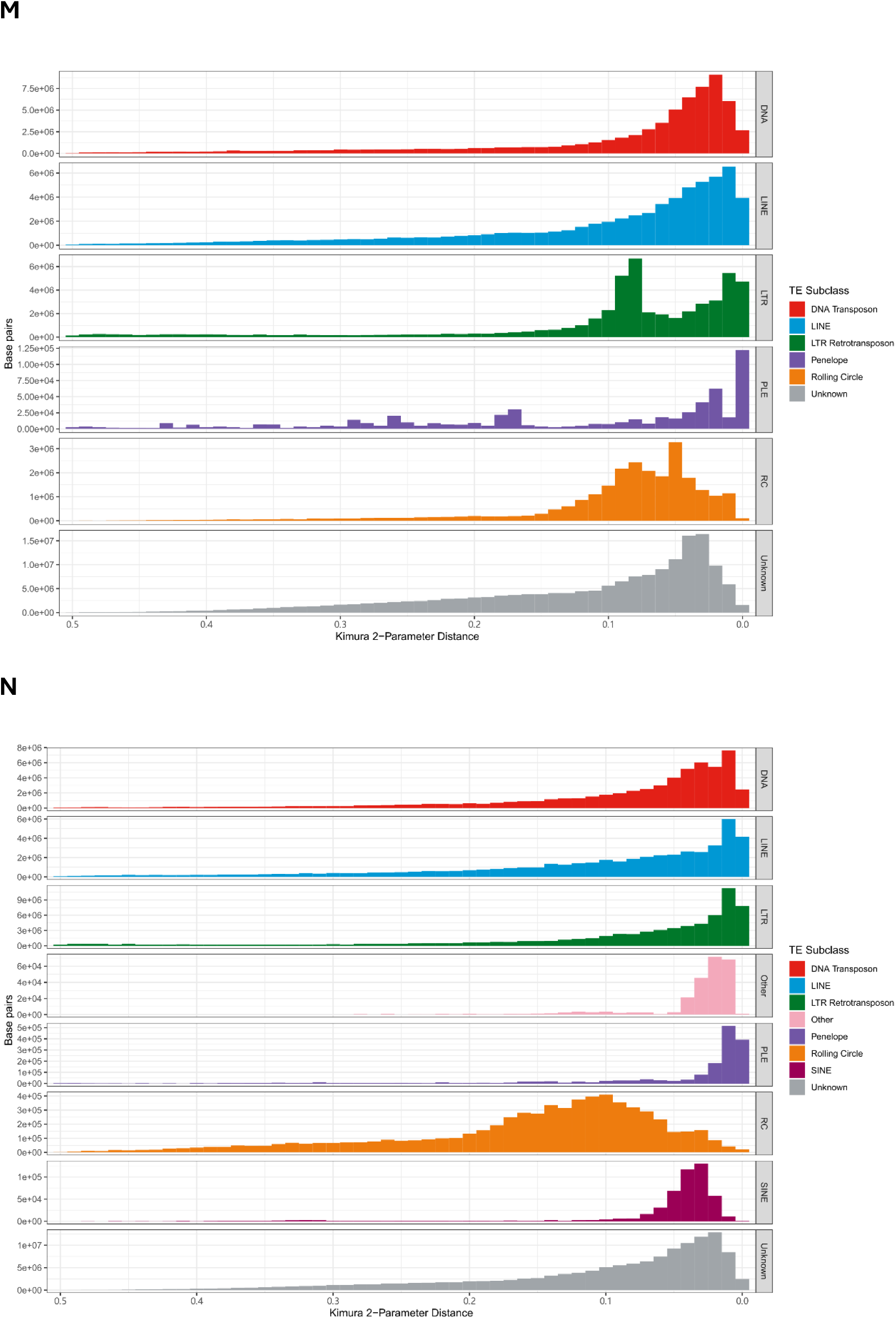
Repetitive elements landscape of: Leptogaster cylindrica (**A**); Neoitamus cyanurus (**B**); Eutolmus rufibarbis (**C**); Machimus rusticus (**D**); Tolmerus cingulatus (**E**); Machimus atricapillus (**F**); Dioctria linearis (**G**); Dioctria rufipes (**H**); Stratiomys singularior (**I**); Microchrysa polita (**J**); Hermetia illucens (**K**); Chorisops tibialis (**L**); Beris morrisii (**M**); Beris chalybata (**N**). The Kimura 2-Parameter distance (X axis) measures the distance between identified repetitive elements and the consensus sequence. A lower Kimura 2-Parameter distance indicates more recent activity of certain repetitive elements. The abundance of each subclass of repetitive elements is measured by the number of its total base pairs.

## Notes

### Competing Interest Statement

The authors have declared no competing interest.

